# Particle Tracking for Quantized Intensities Accelerated with Parallelism

**DOI:** 10.1101/2025.05.30.657103

**Authors:** Lance W.Q. Xu, Nathan Ronceray, Marianna Fanouria Mitsioni, Titouan Brossy, David Šťastný, Radek Šachl, Martin Hof, Aleksandra Radenovic, Steve Pressé

**Affiliations:** Center for Biological Physics, Arizona State University, Tempe, AZ, USA; Department of Physics, Arizona State University, Tempe, AZ, USA; Institute of Bioengineering, School of Engineering, EPFL, Lausanne, Switzerland; NCCR Bio-Inspired Materials, EPFL, Lausanne, Switzerland; J. Heyrovský Institute of Physical Chemistry, Dolejškova 2155/3, CZ-18223 Prague 8, Czech Republic; Faculty of Science, Charles University, Prague, Czech Republic; School of Molecular Sciences, Arizona State University, Tempe, AZ, USA

## Abstract

Traditional particle tracking methods approximate each pixel as reporting a continuous grayscale intensity, an assumption valid for long exposures with many detected photons. At short exposures, however, pixels record only a few discrete photon counts, and this quantization introduces both statistical and computational challenges. We present QTrack, a likelihood-based and parallelized framework that directly models discrete photon detections while exploiting correlations across all frames. By doing so, QTrack surpasses the Cramér – Rao lower bound (CRLB) prediction assuming continuous intensity and localization-and-linking. To make this possible in practice, QTrack exploits the parallelism inherent in both likelihood evaluation and posterior sampling through vectorization, multi-threading, and GPU acceleration, with lightweight inter-thread communication. On a single mid-range GPU (Nvidia GTX 1060, 6 GB), QTrack achieves up to a 50-fold speedup compared to a serial CPU implementation, while maintaining full accuracy and data efficiency. Together, these advances establish that short exposures with quantized photon detections are not a limitation but an opportunity: when modeled rigorously, they enable localization and diffusion coefficient estimates beyond the CRLB prediction for continuous-intensity data, setting a new standard for single-particle tracking.

## 1 Introduction

In widefield single-particle tracking (SPT), camera pixel values are often treated as continuous intensities, an approximation that holds when exposures are long and photon counts are high. At short exposures, however, pixels record only a few photons per frame, and quantization effects become significant. The most extreme case is presented by single-photon avalanche diode (SPAD) arrays^1^, where each pixel reports only a binary outcome (0 or 1). Short exposures and quantized readouts are not simply a limitation, but often a necessity: they provide higher temporal resolution, reduce motion blur for fast-diffusing molecules, and in some applications, discrete photon detections are indispensable—for example, when extracting fluorescence lifetimes from photon arrival times^1–8^ or when separating photons into spectral channels. In this regime, traditional continuous-intensity models of particle tracking break down, and new statistical and computational challenges arise.

Most existing approaches, however, are based on the continuous-intensity assumption. Approximation-based pipelines, most notably localization-and-linking methods^9–11^, detect particle positions frame by frame and then connect them over time. These methods implicitly assume that per-frame photon counts are sufficiently high for localization, which is precisely the assumption underlying the Cramér – Rao lower bound (CRLB) for localization precision^12–16^. As a result, their best possible performance is capped by this CRLB, and they fail in the short-exposure, low-photon regime. Deep learning approaches have been developed under the same assumptions. While they offer computational speed and flexibility^17–25^, their accuracy degrades sharply in low-photon conditions^21^.

In contrast, physics-inspired likelihood-based methods can, in principle, directly model the discrete, quantized photon counts underlying experimental image frames. By leveraging all available photons across all frames, these approaches hold the potential to overturn the long-standing belief that longer exposures and higher per-frame photon counts are always necessary for accurate tracking. Their effectiveness is reflected in strong performance across a wide range of biophysical applications^26–42^. Yet applying likelihood-based methods to SPT remains hampered by one major limitation: computational cost. Evaluating per-pixel likelihoods across particles, frames, and trajectories is intensive, and posterior inference often relies on iterative sampling methods such as MCMC^43^ (see Figs. 1b to 1d), which can require hours of computation on standard CPUs^44^.

**Figure 1:**
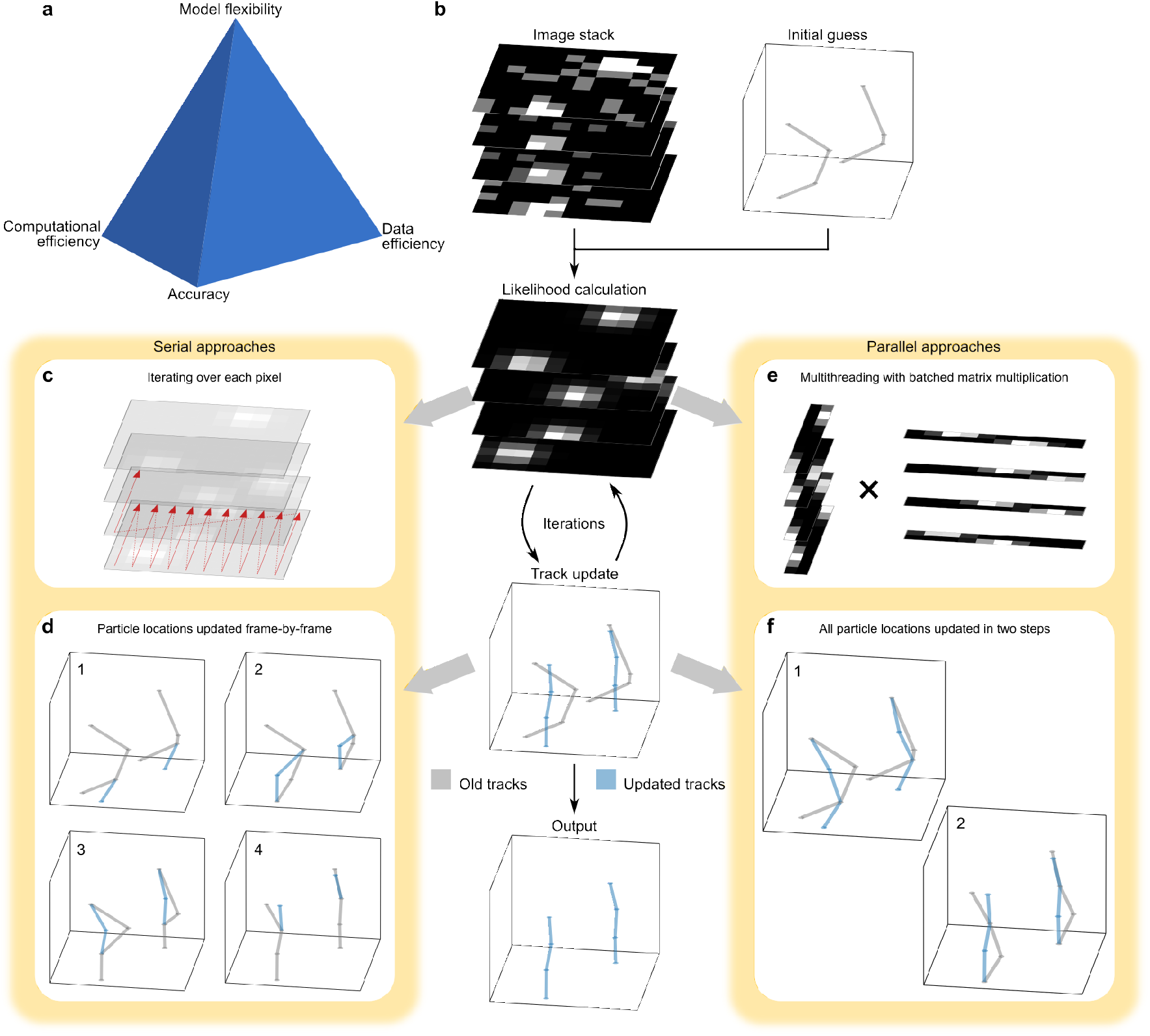
**a**, Particle tracking methods face trade-offs among four key aspects: tracking accuracy, computational efficiency, data efficiency, and model flexibility. These challenges are amplified under short exposures with quantized photon counts, where only a few photons per frame must be used efficiently. **b**, In likelihood-based approaches, tracking begins with a recorded image stack and initial track estimates, followed by iterative updates of particle tracks. Each update requires evaluating the likelihood of the observed data given a proposed set of tracks. **c**, Likelihood calculations can be performed serially by evaluating each pixel individually. **d**, A naive approach to updating tracks involves frame-by-frame serial updates. **e**, Using batched matrix multiplication allows for parallelized likelihood computation, which can more efficiently leverage thousands of threads available in modern GPUs. **f**, With improved algorithmic design, particle positions at even and odd frames can be updated in just two steps, regardless of how many frames there are in total.

Parallelization has been widely explored as a remedy^45–56^, most often through data partitioning or replica-based strategies. However, both approaches break down when strong temporal correlations exist, as in particle tracking: per-pixel likelihoods must be aggregated across frames, and correlations between time points demand frequent synchronization. The resulting communication overhead becomes the dominant bottleneck. *Post hoc* aggregation of subposterior samples^50,52^ introduces further approximations, undermining the statistical rigor that likelihood-based methods are meant to preserve.

To overcome these barriers, we present Quantized Track (QTrack), a likelihood-based SPT framework that explicitly models quantized photon detections. By doing so, QTrack surpasses the localization precision predicted by the CRLB under continuous-intensity assumptions. For example, at an exposure of 10 ms with a diffusion coefficient of 1 µm^2^/s, QTrack improves localization precision by nearly 58%. When the exposure is reduced to 10 µs, where conventional localization-and-linking methods can no longer perform per-frame localization, QTrack outperforms the CRLB prediction by over an order of magnitude. These results establish that short exposures with quantized photon counts are not a limitation but, when properly modeled, a fundamental advantage.

In addition to this conceptual advance, QTrack also addresses the computational bottlenecks that have historically limited likelihood-based approaches. The framework exploits the inherent parallelism of likelihood evaluation and posterior sampling through vectorization, multi-threading, and GPU acceleration, while supporting lightweight inter-thread communication (see Figs. 1e and 1f). On a mid-range GPU (Nvidia GTX 1060, 6 GB), QTrack achieves up to a 50-fold speedup relative to its serial CPU predecessor (Intel i7-7700K), with no loss of accuracy or data efficiency, making high-throughput likelihood-based inference practical.

## 2 Results

We first show that QTrack can surpass the CRLB predictions that assume continuous intensities and a localization-and-linking framework, in both simulated and experimental datasets. By explicitly modeling quantized photon detections, QTrack achieves localization accuracy and diffusion coefficient precision beyond these theoretical limits. Having established this conceptual advance, we then assess the computational performance of QTrack. Compared to its predecessor^44^, QTrack retains the same high level of tracking accuracy, particularly under low-SNR conditions, while delivering a substantial computational boost. On a mid-range Nvidia GTX 1060 (6 GB) GPU, QTrack achieves nearly a 50-fold speedup relative to the original serial implementation on an Intel i7-7700K CPU. We further benchmark computational scaling across parameter regimes, and in the parametric setting—when the number of emitting particles is fixed—the parallel framework achieves an additional 20-fold acceleration, bringing the total speedup to nearly 200-fold on a single GTX 1060 GPU.

### 2.1 QTrack beats the CRLB prediction

We first show, using both simulations and experimental data, that QTrack, by directly modeling discrete photon detections and exploiting spatiotemporal correlations, can surpass the CRLB prediction assuming continuous intensity and localization-and-linking^14^.

Before presenting results, it is helpful to clarify why surpassing this CRLB prediction is possible. In the conventional localization-and-linking paradigm, tracking is performed sequentially: particles are first localized in each frame (implicitly assuming smooth, continuous intensities), and these localizations are then linked across time. The localization step requires a sufficient number of detected photons to achieve precise position estimates^12–16^. With very short exposures, however, only a few discrete photon detections are recorded per frame, leading to large localization errors. Conversely, increasing the exposure time yields more photons but also introduces motion blur, which degrades localization accuracy. Crucially, motion blur in SPT is not limited to translational movement in the imaging plane; it also arises from out-of-plane motion, rotational diffusion, and even sample or system drift. A likelihood-based tracking method such as QTrack, which operates directly on discrete, quantized photon detections while exploiting correlations across frames, avoids this trade-off and can therefore exceed the CRLB prediction assuming continuous intensity and localization-and-linking.

To illustrate this principle, we simulated data with 10 µs exposures. At such short exposures, a particle emitting approximately 10,000 detected photons per second produces, on average, only one detected photon every ten frames. This firmly places the analysis in a regime where photon arrivals must be treated as discrete events. By summing arbitrary numbers of these short-exposure frames, we can effectively construct datasets corresponding to longer exposures, enabling direct comparisons across conditions.

Figure 2a shows particle trajectories inferred by QTrack at 10 ms and 10 µs exposures, while Fig. 2b compares the dynamic localization errors^14^ predicted by the CRLB assuming continuous intensity and localization-and-linking with those obtained using QTrack. Remarkably, even at 10 µs exposure, QTrack yields more accurate position estimates than the CRLB prediction for data acquired with 10 ms exposures. Extending this analysis, Fig. 2c summarizes results across exposures ranging from 10 µs to 30 ms and diffusion coefficients spanning 0.1 µm^2^/s to 10 µm^2^/s, consistently showing that QTrack outperforms the CRLB prediction assuming continuous intensity and localization-and-linking under all tested conditions. For instance, at a diffusion coefficient of 1 µm^2^/s and 10 ms exposure, where the continuous-intensity assumption typically holds, QTrack achieves a dynamic localization error of about 38 nm, compared to the CRLB prediction of 56 nm. When the exposure is reduced to 10 µs, where most frames contain only 0 or 1 photon detection, QTrack’s error increases to about 87 nm, still well below the diffraction limit of 250 nm. In contrast, the CRLB prediction at this exposure rises to 1.13 µm, clearly illustrating the breakdown of the continuous-intensity assumption.

**Figure 2:**
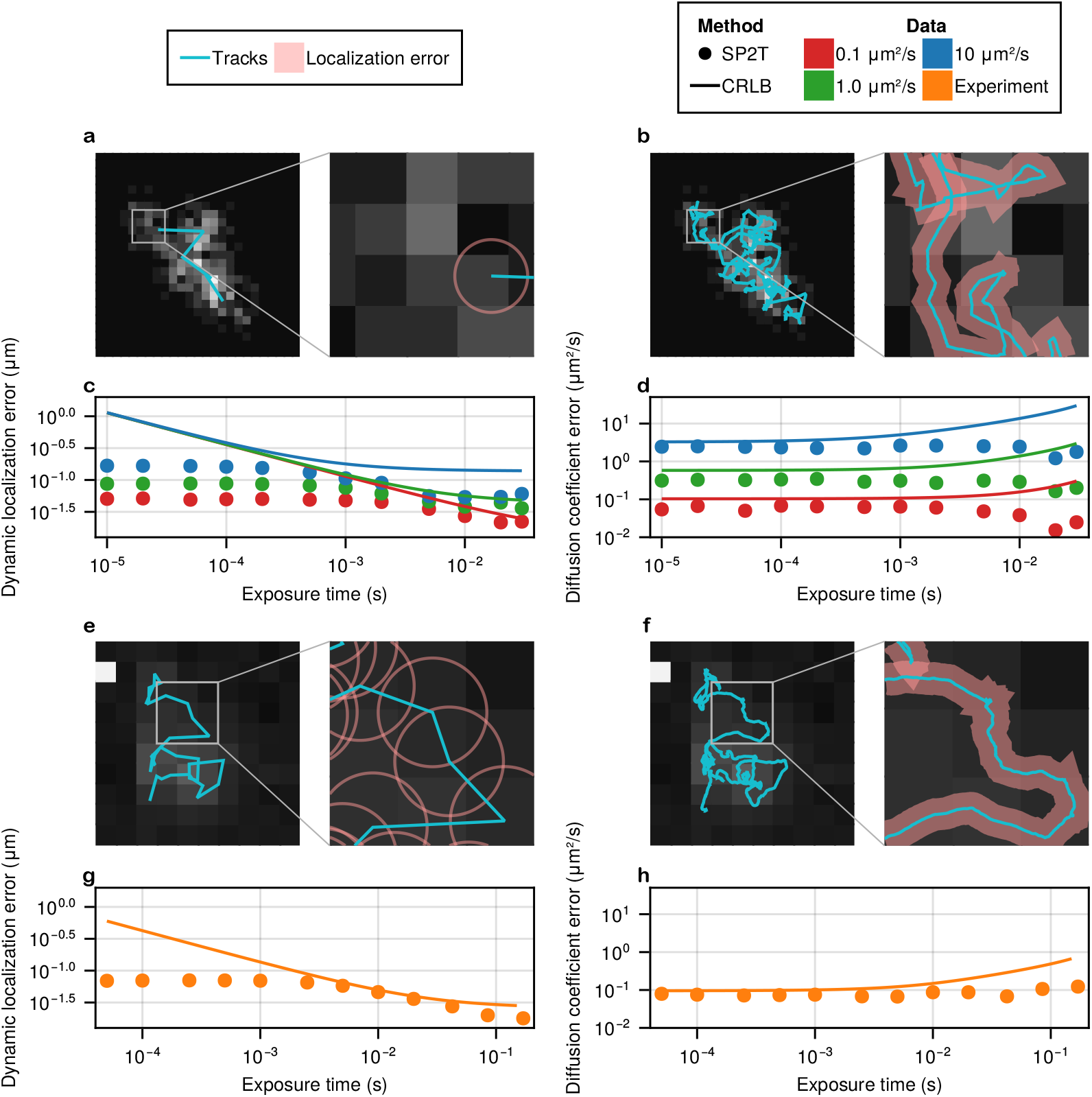
QTrack surpassing the CRLB predictions that assume continuous intensities and a localization-and-linking framework is demonstrated with both simulated and experimental datasets. **a**, Left: particle trajectory from the 10.0 µm^2^/s diffusion simulation with 10 ms exposure, overlaid on the summed image of all frames. Simulation parameters: 10,000 detected photons/s, 590 nm emission wavelength (mimicking Atto565), numerical aperture 1.49, pixel size 100 nm. No background was included to match CRLB conditions^13,14^. Right: zoom-in of the boxed region; the circle indicates the CRLB localization error, which reflects the dynamic localization error (width of the 68% credible interval, CI, accounting for motion blur^14^). **b**, Same as **a**, but for the particle trajectory with 10 µs exposure, where the shaded region indicates localization uncertainty. **c**, Dynamic localization errors from QTrack compared with CRLB predictions across exposures for simulated particles diffusing at 0.1 µm^2^/s, 1.0 µm^2^/s, and 10.0 µm^2^/s. **d**, Uncertainty in diffusion coefficient estimates, given as 68% CI widths, for QTrack versus CRLB predictions using the same datasets. **e–h**, Same layout as **a–d**, but applied to experimental trajectories of Atto565-labeled ganglioside molecules^57,58^. The measured photon budget is about 7,000 photons/s.

The trade-off between localization accuracy and exposure time is not the only limitation of the localization-and-linking scheme. A second trade-off arises when estimating diffusion dynamics. Shorter exposures increase the number of trajectory steps and should, in principle, yield more precise diffusion coefficient estimates. Accordingly, a CRLB prediction also exists for the uncertainty in diffusion coefficient estimation. In Fig. 2d, we compare diffusion coefficient estimates from QTrack with the CRLB predictions assuming continuous intensity and localization-and-linking across a range of exposures and diffusion coefficients, analogous to the localization analysis in Fig. 2e. Once again, QTrack consistently outperforms the CRLB prediction.

Finally, in Figs. 2e to 2h, we repeat the analysis using an experimental dataset of diffusing Atto565-labeled ganglioside molecules^57,58^. The shortest-exposure frames were acquired with the SPAD512^2^ detector from Pi Imaging at 50 µs exposure. Unlike simulations, this real dataset introduces additional complications such as hot pixels, and the total photon budget is reduced to approximately 7,000 photons per second, making it a more challenging case for analysis. Despite these limitations, QTrack consistently outperforms the CRLB prediction assuming continuous intensity and localization-and-linking in both localization accuracy and diffusion coefficient estimation. Specifically, at 10 ms exposure, QTrack achieves a dynamic localization error of 46 nm, compared to the CRLB prediction of 50 nm. When the exposure is reduced to 50 µs, QTrack’s error increases to 69 nm, yet the CRLB prediction at this regime rises sharply to 0.60 µm, highlighting the breakdown of the continuous-intensity assumption and the robustness of QTrack under extreme photon-limited conditions.

### 2.2 Benchmarking QTrack against BNP-Track in the high-photon regime

To confirm that QTrack’s advantages are not limited to low-photon regimes, we next verify that it performs on par with another likelihood-based method optimized for the high-photon count regime, BNP-Track^44^, while offering substantially lower computational cost. This result is particularly notable because quantizing photon detections should, in principle, increase computational expense, as each individual photon arrival must be modeled explicitly.

To test this, we simulated a series of image stacks with identical parameters—particle trajectories, field of view (FOV), pixel size, emission wavelength, numerical aperture, refractive index, and emission rate—varying only the background level. Each stack contained ten 128×128 frames with ten diffusing particles. For illustration, Fig. 3a shows the corresponding time-averaged frames, where the noise level in the rightmost dataset is 20-fold higher than in the leftmost. We then compared QTrack, BNP-Track, and TrackMate^10^, a widely used localization-and-linking tool^59,60,0,62,63^. To ensure a fair comparison, parameters for each method were tuned so that no detections lay more than one diffraction limit from ground truth (spurious detections). Tracking accuracy was then quantified using the detection ratio: the number of correct detections divided by the total number of ground truth positions. As shown in Fig. 3b, both the original and parallel physics-inspired frameworks (BNP-Track and QTrack) reliably detect all particles across all frames, whereas TrackMate’ s performance degrades substantially with increasing background noise.

**Figure 3:**
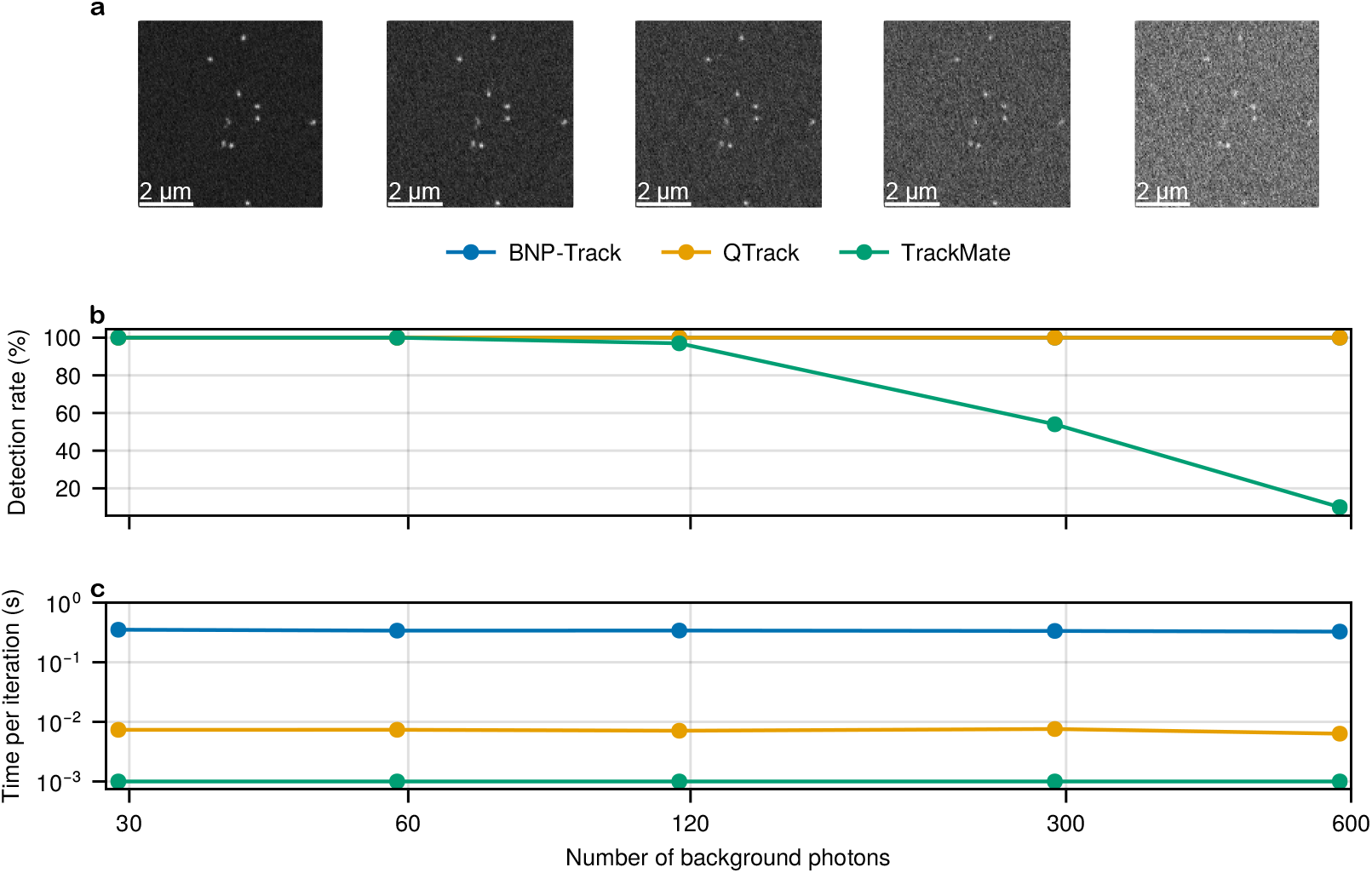
Tracking accuracy and computational efficiency in the high-photon regime, comparing QTrack (likelihood-based), BNP-Track^44^ (likelihood-based), and TrackMate^10^ (localization-and-linking) under varying background noise levels using synthetic data. **a**, Time-averaged frames at different background noise levels, expressed as background photons per pixel per frame, with contrast adjusted for visualization. Each particle contributes about 10,000 photons/s, consistent with the brightness of a real dye^44,64^. **b**, Detection ratio for each method, defined as the fraction of true particle detections without spurious tracks. The QTrack and BNP-Track curves overlap, indicating that their detection ratios are identical across all conditions. **c**, Per-iteration runtime for each method. Simulation parameters: numerical aperture 1.45, refractive index 1.515, emission wavelength 665 nm, frame exposure 33 ms, and pixel size 133 nm. Each dataset contained ten 128×128 frames with particles diffusing at 0.1 µm^2^/s. Benchmarks were run on the same desktop computer (Intel i7-7700K CPU, Nvidia GTX 1060, 6 GB GPU); see Computational efficiency measurement.

Having evaluated tracking accuracy using the detection ratio, we now turn to compu-tational efficiency. Unlike accuracy metrics, runtime comparisons across methods are more nuanced because different algorithms infer different quantities from the same data. For example, QTrack and BNP-Track (both physics-inspired likelihood-based methods) estimate full particle trajectories along with the uncertainty of each track, capabilities absent in localization-and-linking tools such as TrackMate.

Practical efficiency is also shaped by how much manual parameter tuning is required. TrackMate demands user-specified parameters such as localization quality thresholds and maximum linking distances, which are often optimized through trial-and-error. This process can be time-consuming, dataset-dependent, and highly variable across users. By contrast, likelihood-based methods involve relatively few tunable hyperparameters^30,43,44,65,66^, and are typically designed to run in a fully automated manner once the raw data are provided^30,42,65,67–71^. Thus, for likelihood-based frameworks, the “user time” cost is negligible, and runtime reflects only the computation itself.

Despite these caveats, comparing per-iteration runtime remains informative for highlighting algorithmic and implementation differences. This comparison is straightforward between BNP-Track and QTrack: both rely on MCMC sampling, follow the same inference logic, and require a similar number of iterations to achieve equivalent accuracy. The key distinction is that QTrack models quantized photon detections—an operation that should, in principle, be more expensive—yet runs dramatically faster thanks to its parallelization scheme. For TrackMate, we define per-iteration runtime as the time required for one round of localization and linking with fixed parameters, excluding manual tuning, to provide a fairer algorithmic baseline.

As shown in Fig. 3c, all three methods exhibit similar per-iteration times across signal-to-noise ratio (SNR) regimes, consistent with the fact that SNR primarily affects the number of iterations required for convergence rather than the cost per iteration. TrackMate reports the lowest per-iteration time (typically under 10 ms on an Intel i7-7700K CPU, 4 cores, 8 threads, up to 4.5 GHz), but this excludes substantial user tuning time. BNP-Track and QTrack, in contrast, are fully automated. Critically, QTrack achieves nearly a 50-fold reduction in computational cost on a single Nvidia GTX 1060 (6 GB) GPU compared to BNP-Track running on the same CPU, underscoring how parallelization makes rigorous, quantized-likelihood inference both practical and scalable.

In addition to the tests shown in Fig. 3, we conducted a complementary experiment in which the brightness of each emitting particle varied across datasets. The results, shown in Fig. 4, are consistent with those in Fig. 3, further confirming that the parallel physics-inspired framework maintains its performance while being nearly 50 times more computationally efficient.

**Figure 4:**
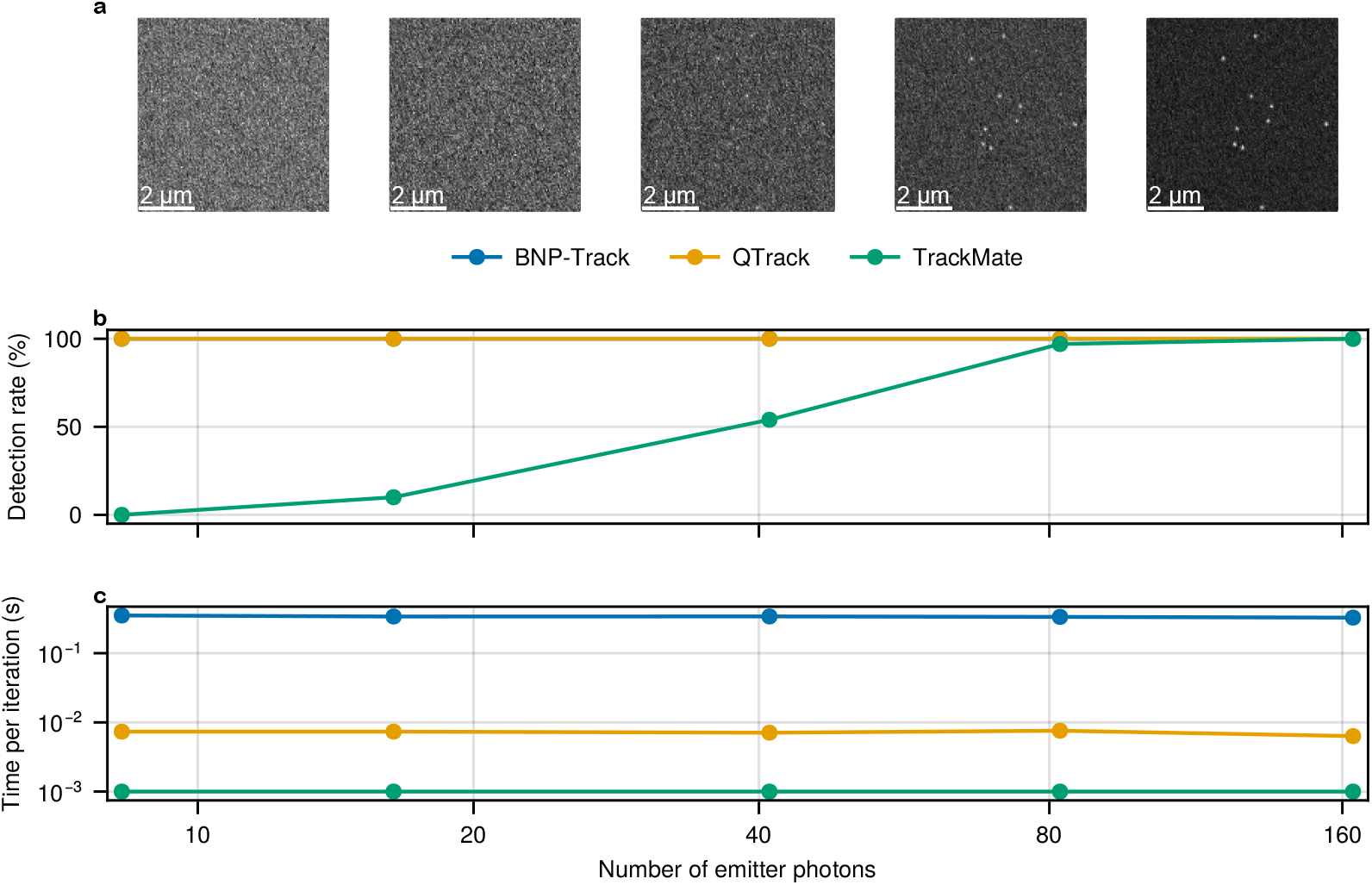
Tracking accuracy and computational efficiency in the high-photon regime, comparing QTrack (likelihood-based), BNP-Track^44^ (likelihood-based), and TrackMate^10^ (localization-and-linking) under varying particle emission rates. **a**, Time-averaged frames at different emission rates; background is fixed at about 60 photons per pixel per frame. **b**, Detection ratio for each method, defined as the fraction of true particle detections without spurious results. QTrack (orange) and BNP-Track (blue) achieve identical detection ratios across all emission rates, while TrackMate degrades at lower photon counts. **c**, Per-iteration runtime for each method, illustrating computational efficiency. Simulation parameters: numerical aperture 1.45, refractive index 1.515, emission wavelength 665 nm, frame exposure 33 ms, pixel size 133 nm. Each dataset contained ten 128×128 frames with particles diffusing at 0.1 µm^2^/s. All benchmarks were performed on the same desktop computer (Intel i7-7700K CPU, Nvidia GTX 1060, 6 GB GPU); see Computational efficiency measurement.

### 2.3 Computational efficiency scaling

Building on the previous subsection, we now show that QTrack continues to outperform BNP-Track even under more demanding parameter variations, despite explicitly modeling quantized photon detections. We provide detailed benchmarks of how both methods scale with frame size, number of frames, and number of emitting particles. For a comprehensive assessment across hardware platforms, we evaluate QTrack on both GPU and CPU, alongside BNP-Track on CPU.

In Fig. 5a, we simulate 100 frames, each containing two emitting particles, with frame sizes ranging from 16×16 to 1024×1024 pixels. QTrack consistently maintains a tenfold speed advantage over BNP-Track on the CPU, with scaling efficiency improving as frame size increases. This gain reflects the optimized algorithmic structure described in Methods. On the GPU, QTrack delivers an additional fourfold speedup for frames of size 32×32 and larger. For 16×16 frames, however, GPU performance matches that of the CPU because the per-kernel workload is too small, and communication overhead between the CPU and GPU dominates^72,73^.

**Figure 5:**
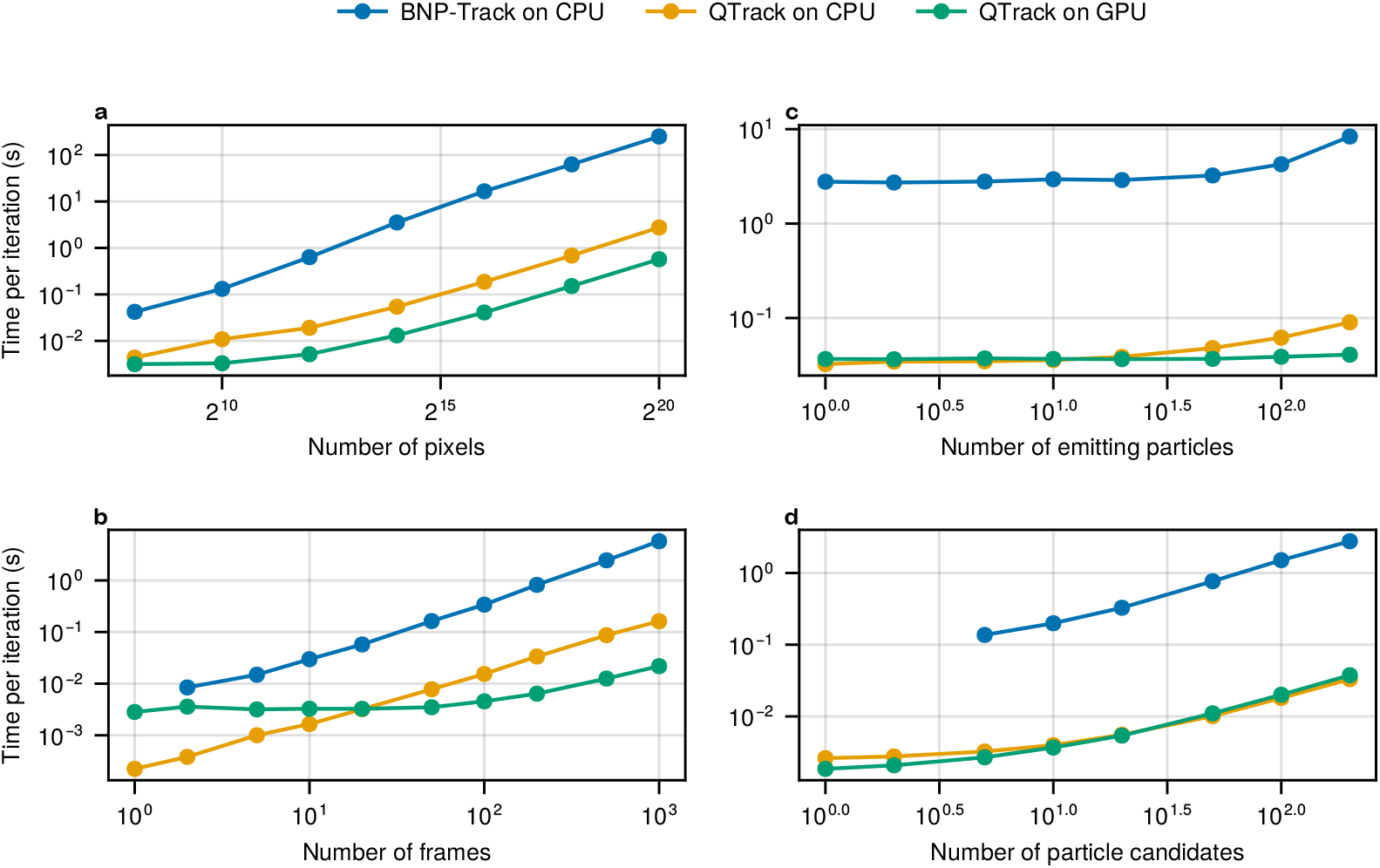
Scalability of computational efficiency comparing BNP-Track (CPU, blue) with QTrack on CPU (orange) and QTrack on GPU (green), evaluated across key parameters. **a**, Per-iteration runtime versus the number of pixels, from 16×16 up to 1024×1024. **b**, Runtime versus the number of frames, from 1 to 1000. **c**, Runtime versus the number of emitting particles, from 1 to 200. **d**, Runtime versus the number of particle candidates (the maximum number of particles allowed in the FOV, set by the user), also from 1 to 200. Simulation parameters: numerical aperture 1.45, refractive index 1.515, emission wavelength 665 nm, frame exposure 33 ms, and pixel size 133 nm. Each dataset contained ten 128×128 frames with particles diffusing at 0.1 µm^2^/s. Benchmarks were run on the same desktop computer (Intel i7-7700K CPU, Nvidia GTX 1060, 6 GB GPU); see Computational efficiency measurement.

Figure 5b examines how per-iteration time scales with the number of frames analyzed. Here, we simulated two emitting particles across 1000 frames of size 50×50 and then analyzed subsets ranging from a single frame to the full stack. Because of its design, BNP-Track cannot process single-frame datasets, making it unsuitable for localization tasks. Consistent with earlier results, QTrack achieves *∼*20-fold speedups on the CPU. On the GPU, however, a computational advantage only emerges for datasets with more than 20 frames; for smaller datasets, CPU – GPU communication overhead outweighs the benefits of parallelism.

We next evaluate how per-iteration time scales with the number of emitting particles. Beginning with ten 128×128 frames containing a single particle, we incrementally increased the number of emitters up to 200. As shown in Fig. 5c, both BNP-Track and QTrack on the CPU remain relatively stable up to about 20 particles, after which computational cost grows more noticeably. By contrast, QTrack on the GPU remains largely unaffected even as the number of emitting particles increases, highlighting its efficiency in high-multiplicity regimes.

This scaling behavior is linked to the nonparametric structures underlying physics-inspired frameworks^43,74–76^. Both QTrack and BNP-Track assume a fixed, user-specified number of candidate particles, only a subset of which are active emitters. The remainder serve as “invisible” placeholders. Each candidate carries a binary indicator determining whether it is active, and these indicators are updated for all candidates at every iteration. As a result, computational cost scales with the total number of candidate particles, not just the active ones.

The advantages of combining nonparametric posterior structures with efficient sampling strategies, central to QTrack, are highlighted in Fig. 5d. Here, we fix the number of emitters to one while increasing the candidate pool from 1 to 200. As expected, scaling becomes more regular, reflecting the framework’s algorithmic design. Two points stand out: first, BNP-Track does not permit fewer than five candidate particles, limiting flexibility in this regime. Second, with only one true emitter, QTrack shows minimal differences between CPU and GPU performance, underscoring that GPU parallelism provides the greatest benefit when the number of emitting particles is large.

### 2.4 Nonparametric cost

Having established the importance of nonparametric structures in physics-inspired frameworks and their role in increasing the complexity of posterior sampling, we next quantify the computational cost of this modeling flexibility. To do so, we compare two configurations of QTrack: a nonparametric mode with 200 candidate particles (as in earlier sections), and a parametric mode in which the number of candidates is fixed to the true number of emitters. Both modes are evaluated on CPU and GPU to assess hardware-dependent performance. As shown in Fig. 6, the parametric mode is consistently more efficient across platforms. In particular, for datasets with 200 emitters, the GPU-accelerated parametric version achieves an additional 20-fold speedup relative to its nonparametric counterpart.

**Figure 6:**
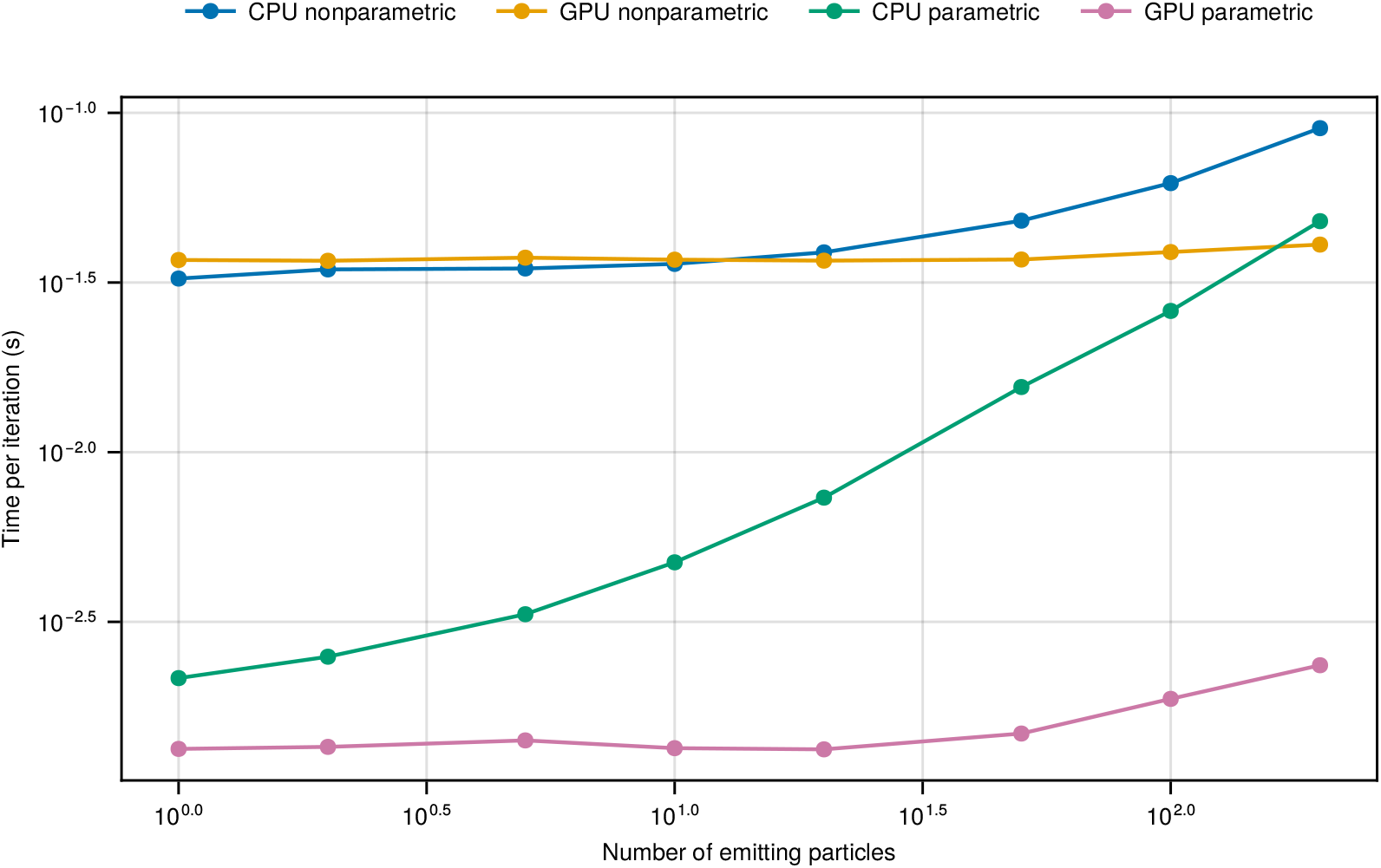
Trade-off between nonparametric flexibility and computational efficiency in QTrack, benchmarked on both CPU and GPU. We compare QTrack in its nonparametric mode (200 candidate particles, with the number of true emitters ranging from 1 to 200) against its parametric mode (number of candidates fixed to the true number of emitters). All benchmarks were performed on the same desktop computer with an Intel i7-7700K CPU and Nvidia GTX 1060 (6 GB) GPU; see Computational efficiency measurement.

These findings underscore a fundamental trade-off for likelihood-based methods: non-parametric models provide flexibility and robustness in scenarios where the number of emitters is unknown or time-varying, but at a higher computational cost. The decision between parametric and nonparametric formulations should therefore be guided by application-specific requirements, balancing modeling generality against available computational resources.

## 3 Discussion

To tackle particle tracking in the short-exposure, low-photon-count regime, developing a physics-inspired, likelihood-based framework that directly models quantized photon detections and spatiotemporal correlations becomes essential. Here, we introduce QTrack, a tracking framework designed explicitly for this purpose. Conventional localization-and-linking schemes, constrained by photon counts per frame and motion blur, are unable to reach the same level of accuracy. In contrast, QTrack consistently surpasses these theoretical performance limits. In the high – photon-count regime, QTrack improves localization precision by over 50% relative to the CRLB prediction assuming continuous intensity and localization-and-linking. In the low – photon-count, quantized regime, QTrack achieves even larger gains, delivering more than an order-of-magnitude improvement in localization precision.

The main cost of integrating such realistic physics into data analysis is the substantial increase in computational demand. Traditional embarrassingly parallel strategies distribute workloads across large CPU clusters^45–51,54,55^, but these approaches are most effective only when inference tasks require little communication between processes. This assumption breaks down in systems governed by tightly coupled physical constraints, such as time-correlated motion in particle tracking, where frequent communication between frames or model components is unavoidable. In such cases, accelerating a single inference chain remains the critical bottleneck.

To overcome this challenge, QTrack employs a fundamentally different parallelization strategy. Instead of distributing independent chains or partitioning data across nodes, QTrack exploits the fine-grained parallelism inherent in its likelihood structure, parallelizing both likelihood evaluations and sampling updates within a single inference chain. By leveraging frequent yet lightweight communication between CPU and GPU, QTrack is optimized for modern desktop-class hardware. This design makes high-throughput Bayesian inference feasible without requiring large-scale computing infrastructure, while still preserving the accuracy and data efficiency of BNP-Track. Importantly, QTrack achieves up to a 50-fold speedup on a mid-range GPU, transforming it from a powerful but time-intensive tool into one capable of practical, high-throughput single-particle tracking.

At the same time, QTrack’s design does not preclude further acceleration through conventional parallelization strategies. As it does not rely on embarrassingly parallel schemes, QTrack can be combined with them, for example, by running multiple inference chains or distributing datasets across compute nodes, to extend its efficiency beyond the desktop.

In addition to hardware advances (*e*.*g*., TPUs for enhanced performance), several emerging computational techniques could further accelerate QTrack. We highlight three particularly promising directions: (1) Mixed-precision computing^77–79^, which allows QTrack to represent different variables at different precision levels—for example, storing object positions at lower precision while maintaining probability density estimates at higher precision —thereby reducing memory usage and improving runtime without sacrificing accuracy; (2) Randomized numerical linear algebra^80^, which leverages stochastic approximations for large-scale matrix operations, offering scalability in high-dimensional tracking problems; (3) Deep learning – based acceleration^56,81,82^, where architectures such as normalizing flows^83^ or diffusion models^84,85^ could approximate computational bottlenecks in physics-inspired inference, either by learning surrogates for complex likelihoods or proposing efficient posterior samples^81^. In parallel, advances in SPAD detector hardware, such as higher photon-detection efficiency, fewer dead pixels per array, and improved timing jitter, will directly enhance SNR and temporal fidelity. These detector-side improvements are orthogonal to QTrack’s inference scheme and thus complement, rather than alter, our conclusions.

By providing concrete strategies for principled, non-naïve parallelization that preserve mathematical rigor and statistical accuracy, QTrack demonstrates that likelihood-based inference can keep pace with modern computational demands. Through techniques such as vectorization, multi-threading, and GPU acceleration, we are no longer constrained to only small or simple systems, nor limited to problems where the likelihood is embarrassingly parallelizable^45–49,51,54,55^. Moreover, the rise of modern programming languages and libraries^86–90^ lowers the barrier to adopting these acceleration techniques, making them broadly accessible to the scientific community.

Looking forward, we envision a growing role for scalable, interpretable, and parallel likelihood-based frameworks that approach the speed of black-box approximations while retaining transparency and physical grounding. As datasets continue to expand in size and complexity, bridging the gap between statistical rigor and computational scalability will be essential. QTrack represents a concrete step toward this goal—delivering accuracy, efficiency, and flexibility for tracking fast, noisy, and crowded biological systems without compromise.

While demonstrated here for two-dimensional diffusion, the same framework could naturally be extended to axial dynamics by incorporating depth-sensitive imaging, *e*.*g*., multiplane or astigmatic PSFs, into the forward model and inference (see Methods).

## Acknowledgments

S.P. acknowledges support from the NIH (R35GM148237), ARO (W911NF-23-1-0304), and NSF (Grant No. 2310610). N.R. and A.R. acknowledge support from the European Research Council (grant 101020445). M.F.M. acknowledges support from the National Center of Competence in Research Bio-Inspired Materials (NCCR 51NF40-182881). D.Š., R.Š. and M.H. acknowledge the Advanced Multiscale Materials for Key Enabling Technologies project, supported by the Ministry of Education, Youth, and Sports of the Czech Republic, Project No. CZ.02.01.01/00/22_008/0004558, Co-funded by the European Union. D.Š., R.Š., and M.H. acknowledge GAČR grant 25-15346S provided by the Czech Science Foundation.

## Data availability

All data supporting the findings of this study are available as benchmark datasets included with the accompanying code repository.

## Code availability

Our code is publicly available at https://github.com/LabPresse/SP2T.jl.

## Competing interests

The authors declare no competing interests.

## 4 Methods

This section introduces the core parallelization and optimization strategies employed in the proposed parallel framework. For clarity, we define the following notation: let *N* denote the number of frames; *I* and *J* represent the number of pixels in the *x*- and *y*-directions per frame, respectively; and 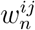 denote the binary measurement collected at pixel (*i, j*) in frame *n*. Let *M* be the number of emitting particles, with 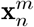 represent-ing the position of particle *m* at frame *n*. The complete dataset can then be written as 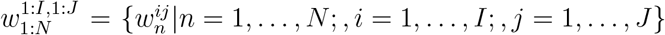, and the full set of particle tracks as 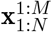.

As discussed in the Introduction, the goal of tracking is to establish a full probability distribution over *M* (the number of particles) and 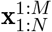 (all elements of all tracks), given 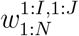 (the recorded frames). In addition, we will also need to estimate the mean squared displacement (MSD) of the particles, as it is typically unknown *a priori*. Using the notations introduced so far, we can express this target probability distribution as 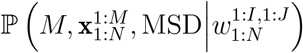.

In the Bayesian paradigm, this target probability distribution is termed the posterior distribution, representing the probability distribution of all variables of interest given the collected data. To construct this posterior, we apply Bayes’ theorem and get

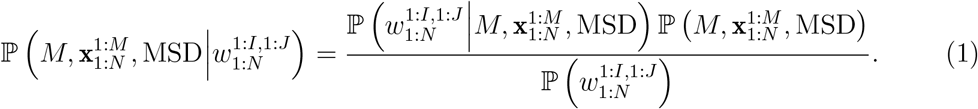

On the right hand side, the first term in the numerator, 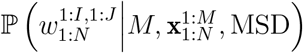, represents the likelihood, which is the probability density of the data given the variables of interest. This term usually describes the system’s physics (the model for photon detection in this context) and realistic complications such as hot pixels. The second term, 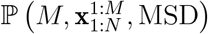, known as the prior, imposes additional constraints on the variables of interest, such as a particle’s motion model and ensuring the MSD remains non-negative. The denominator, 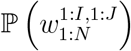, is referred to as the evidence and is generally treated as a normalization constant, as it does not depend on any variable of interest.

Our parallel framework employs Gibbs sampling^43,91^ to infer the variables of interest. This iterative procedure updates each variable by sampling from its conditional posterior distribution, holding the others fixed. Specifically, the particle trajectories 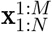 are first updated from the conditional posterior 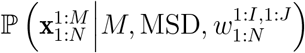, followed by updating the number of emitting particles 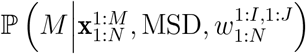, and finally, the mean squared displacement MSD is updated using 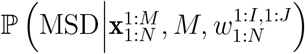.

### 4.1 Likelihood parallelization

As introduced above, the likelihood 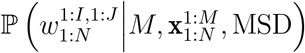 represents the probability distribution of the data given the number of particles, the particle tracks, and the MSD. Since it describes the system’s physics, it is usually the most computationally expensive term in the posterior distribution. Therefore, we first focus on parallelizing this term to improve the overall computation time of the framework.

To express this likelihood mathematically, we first observe that once all particle tracks are known, neither the number of particles nor the MSD affects photon detection. This observation allows us to simplify 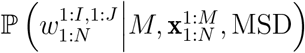 to 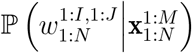.

Furthermore, considering that we are dealing with fluorescently emitting particles whose lifetimes (on the order of nanoseconds^92^) are significantly shorter than the frame period (microseconds and above), the probability that an emitting particle excited during the exposure period of one frame will result in a photon detection in the next frame is negligible. Therefore, it is reasonable to assume that particle positions at frame *n* do not influence other frames. This assumption allows us to further simplify the likelihood as a product:

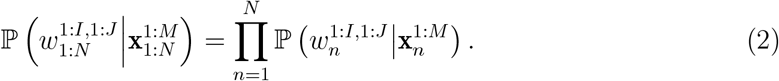

Similarly, since the pixels of a detector usually operate independently, we can express the per-frame likelihood 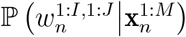 as a product over the individual pixel likelihoods:

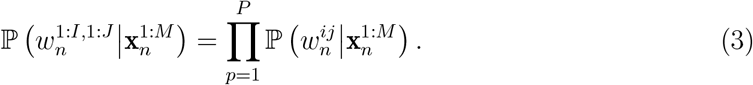

The per-pixel likelihood 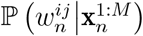 is influenced by four key factors: the point spread function (PSF) of the optical system, the pixel area *A*_*ij*_, the brightness of the particles *h*, and the number of emitting particles *M*. The PSF characterizes how light from a point source is distributed across the detector^16^. At the same time, brightness represents the expected number of photons arriving at a pixel perfectly aligned with the PSF of a particle located at the focal plane. The total number of photons received by pixel *p* from all emitting particles, denoted as 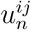, can be expressed as:

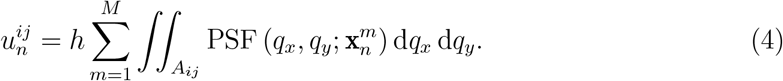

Then 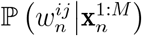 can be constructed using equation (4) and 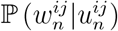, which is specific to the detector and the experimental conditions. In QTrack, photon detections are modeled probabilistically through the detection distribution associated with each pixel. For SPAD-based detectors, the Bernoulli distribution provides a natural model in the low-intensity regime, where the expected photon count 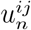 per exposure is so small that the recorded value 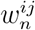 is effectively binary (0 or 1). In this case, each pixel simply records whether a photon was detected during the exposure interval. More generally, however, the correct distribution is Binomial, reflecting the probability of detecting *k* photons out of a finite number of independent trials, each with detection probability 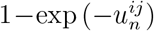 QTrack therefore implements the Binomial distribution as the default model, with the Bernoulli case arising naturally as the low-intensity limit. For completeness, we also account for CCD and sCMOS detectors, which integrate many photon arrivals per pixel; in this case, the probability of readouts 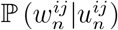 is often approximated by a Gaussian distribution^16^.

In equation (4), the form of the PSF is often approximated as a Gaussian function of^16^:

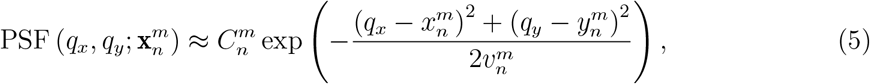

where 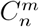 is some normalization constant and 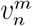 is the variance of the Gaussian function in the lateral direction, which is dependent on the emission wavelength, numerical aperture, and a particle’s *z*-location. With this approximation, equation (4) can be written as a matrix multiplication:

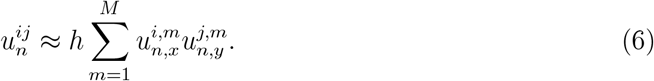

Here, 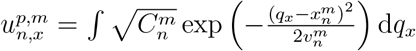 which can be calculated using the error function.

It follows that the expected photon count at all pixels on frame *n* can be expressed as a matrix multiplication:

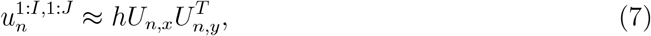

where *U*_*n,x*_ is an *I × M* matrix with 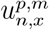 being its elements. To calculate 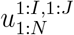, we can perform matrix multiplications for all *N* frames in parallel. This operation is known as batched matrix multiplication. It is implemented in many numerical and deep-learning libraries^88–90,93–96^. Similarly, as all elements in 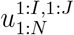 are calculated, each 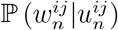 can be calculated in parallel as well using techniques such as broadcasting^97–99^.

### 4.2 Two-step track update

As described above, particle trajectories are updated by sampling from the conditional posterior distribution 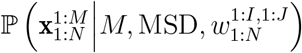 within the Bayesian framework. Based on the concept of Markov boundary^43,100^, the minimal set of random variables containing all useful information to infer another random variable, our parallel framework achieves this through a two-step update scheme designed to maximize parallel efficiency. First, we compute the conditional distribution 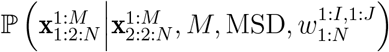, where 1:2:*N* and 2:2:*N* denote all odd and even frame indices, respectively. By holding the particle positions at all even frames fixed, the positions at odd frames become conditionally independent and can thus be updated in parallel. In the second step, we compute the conditional distribution for the even indices, 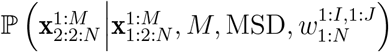, and update all even frame positions in parallel while holding the odd ones fixed. This alternating blockwise update strategy allows all emitting particle tracks to be refreshed in just two passes, each enabling full parallelization across frames.

### 4.3 Memory management

In addition to leveraging parallelism, our framework enhances computational efficiency through careful memory management. A key optimization involves ensuring contiguous memory allocation for the most computationally intensive components of the algorithm^73,101,102^. In particular, the likelihood term, 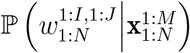, is computed using only the *M* emitting particles. Our parallel framework dynamically reorganizes particle tracks to optimize this step so that the data associated with emitting particles is stored in a contiguous memory block.

Another key memory management strategy employed in our framework is minimizing data copying and avoiding the overhead associated with garbage collection—reclaiming memory no longer in use^101,103^. To achieve this, our framework preallocates memory for intermediate variables—such as 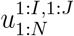 —at the start of execution and reuses this memory throughout the computation.

Similarly, to minimize communication overhead between the CPU and GPU, our framework is designed so that large data transfers occur only once at the beginning of execution. Only lightweight scalar values are exchanged between the CPU and GPU during runtime, significantly reducing latency and maximizing computational throughput.

### 4.4 Computational efficiency measurement

All the tests presented in this study were performed on a desktop computer equipped with an Intel i7-7700K CPU (four cores, eight threads, up to 4.5 GHz), an Nvidia GTX 1060 6 GB GPU, 64 GB of RAM, and Ubuntu 24.04.2 as the operating system. To evaluate the per-iteration runtime of BNP-Track^44^ and the proposed parallel framework, each method was executed through an initial test run to account for and complete any initialization overhead. Subsequently, the runtime for 1000 iterations was measured, and the reported per-iteration time was computed by dividing the total runtime by 1000.

### 4.5 Software

Our framework is implemented in Julia^86^, a high-level, high-performance dynamic programming language designed for technical computing. The framework leverages several core Julia packages: CUDA.jl^87^ for GPU acceleration on Nvidia hardware, Makie.jl^104^ for high-quality data visualization, Distributions.jl^105^ for probabilistic modeling and working with statistical distributions, and Flux.jl^90,93^ for efficiently implementing batched matrix operations and numerical routines.

## Notes

### Competing Interest Statement

The authors have declared no competing interest.

### Summary of Updates

The Results section has been updated with experimental data on ganglioside.

## References

1. Ronceray, N. et al. Wide-field fluorescence lifetime imaging of single molecules with a gated single-photon camera. Light Sci. Appl. 14. issn: 2047-7538 (2025).

2. Ulku, A. C. et al. A 512 × 512 SPAD Image Sensor with Integrated Gating for Widefield FLIM. IEEE J. Sel. Topics Quantum Electron. 25, 1. issn: 1558-4542 (2019).

3. Castello, M. et al. A robust and versatile platform for image scanning microscopy enabling super-resolution FLIM. Nat. Methods 16, 175. issn: 1548-7105 (2019).

4. Smith, J. T. et al. In vitro and in vivo NIR fluorescence lifetime imaging with a time-gated SPAD camera. Optica 9, 532. issn: 2334-2536 (2022).

5. Fazel, M. et al. High Resolution Fluorescence Lifetime Maps from Minimal Photon Counts. ACS Photonics 9, 1015. issn: 2330-4022 (2022).

6. Ronceray, N. et al. Liquid-activated quantum emission from pristine hexagonal boron nitride for nanofluidic sensing. Nat. Mater. 22, 1236. issn: 1476-4660 (2023).

7. Fazel, M. et al. Building Fluorescence Lifetime Maps Photon-by-Photon by Leveraging Spatial Correlations. ACS Photonics 10, 3558. issn: 2330-4022 (2023).

8. Bucci, A. et al. 4D Single-particle tracking with asynchronous read-out single-photon avalanche diode array detector. Nat. Commun. 15, 6188. issn: 2041-1723 (2024).

9. Chenouard, N. et al. Objective comparison of particle tracking methods. Nat. Methods 11, 281. issn: 1548-7105 (2014).

10. Tinevez, J.-Y. et al. TrackMate: An open and extensible platform for single-particle tracking. Methods 115, 80. issn: 1046-2023 (2017).

11. Jaqaman, K. et al. Robust single-particle tracking in live-cell time-lapse sequences. Nat. Methods 5, 695. issn: 1548-7105 (2008).

12. Thompson, R. E., Larson, D. R. & Webb, W. W. Precise Nanometer Localization Analysis for Individual Fluorescent Probes. Biophys. J. 82, 2775. issn: 0006-3495 (2002).

13. Michalet, X. Mean square displacement analysis of single-particle trajectories with localization error: Brownian motion in an isotropic medium. Phys. Rev. E 82, 041914. issn: 1550-2376 (2010).

14. Michalet, X. & Berglund, A. J. Optimal diffusion coefficient estimation in single-particle tracking. Physical Review E 85, 061916. issn: 1550-2376 (2012).

15. Lelek, M. et al. Single-molecule localization microscopy. Nat. Rev. Methods Primers 1, 39. issn: 2662-8449 (2021).

16. Fazel, M. et al. Fluorescence microscopy: A statistics-optics perspective. Rev. Mod. Phys. 96, 025003. issn: 1539-0756 (2 2024).

17. Zhong, Y., Li, C., Zhou, H. & Wang, G. Developing Noise-Resistant Three-Dimensional Single Particle Tracking Using Deep Neural Networks. Anal. Chem. 90, 10748. issn: 1520-6882 (2018).

18. Spilger, R. et al. A Recurrent Neural Network for Particle Tracking in Microscopy Images Using Future Information, Track Hypotheses, and Multiple Detections. IEEE Trans. Image Process. 29, 3681. issn: 1941-0042 (2020).

19. Yao, Y., Smal, I., Grigoriev, I., Akhmanova, A. & Meijering, E. Deep-learning method for data association in particle tracking. Bioinformatics 36 (ed Wren, J.) 4935. issn: 1367-4811 (2020).

20. Spilger, R. et al. Deep probabilistic tracking of particles in fluorescence microscopy images. en. Med. Image Anal. 72, 102128 (2021).

21. Cheng, H.-J., Hsu, C.-H.Hung, C.-L. & Lin, C.-Y. A review for cell and particle tracking on microscopy images using algorithms and deep learning technologies. Biomed. J. 45, 465. issn: 2319-4170 (2022).

22. Qiao, C. et al. Evaluation and development of deep neural networks for image super-resolution in optical microscopy. Nat. Methods 18, 194. issn: 1548-7105 (2021).

23. Song, D. et al. Deep Learning-Assisted Automated Multidimensional Single Particle Tracking in Living Cells. Nano Lett. 24, 3082. issn: 1530-6992 (2024).

24. Qiao, C. et al. A neural network for long-term super-resolution imaging of live cells with reliable confidence quantification. Nat. Biotechnol. issn: 1546-1696 (2025).

25. Qiao, C. et al. Fast-adaptive super-resolution lattice light-sheet microscopy for rapid, long-term, near-isotropic subcellular imaging. Nat. Methods 22, 1059. issn: 1548-7105 (2025).

26. Jazani, S., Sgouralis, I. & Pressé, S. A method for single molecule tracking using a conventional single-focus confocal setup. J. Chem. Phys. 150. issn: 1089-7690 (2019).

27. Jazani, S. et al. An alternative framework for fluorescence correlation spectroscopy. Nat. Commun. 10, 3662. issn: 2041-1723 (2019).

28. Tavakoli, M. et al. Pitching Single-Focus Confocal Data Analysis One Photon at a Time with Bayesian Nonparametrics. Phys. Rev. X 10, 011021. issn: 2160-3308 (2020).

29. Karslake, J. D. et al. SMAUG: Analyzing single-molecule tracks with nonparametric Bayesian statistics. Methods 193, 16. issn: 1046-2023 (2021).

30. Van de Schoot, R. et al. Bayesian statistics and modelling. Nat. Rev. Methods Primers 1. issn: 2662-8449 (2021).

31. Kilic, Z. et al. Extraction of rapid kinetics from smFRET measurements using integrative detectors. Cell Rep. Phys. Sci. 2, 100409. issn: 2666-3864 (2021).

32. Kilic, Z., Sgouralis, I. & Pressé, S. Generalizing HMMs to Continuous Time for Fast Kinetics: Hidden Markov Jump Processes. Biophys. J. 120, 409. issn: 0006-3495 (2021).

33. Vroylandt, H., Goudenège, L., Monmarché, P., Pietrucci, F. & Rotenberg, B. Likelihood-based non-Markovian models from molecular dynamics. Proc. Natl. Acad. Sci. U.S.A. 119. issn: 1091-6490 (2022).

34. Wu, Y.-L. et al. Maximum-likelihood model fitting for quantitative analysis of SMLM data. Nat. Methods 20, 139. issn: 1548-7105 (2022).

35. Linka, K. et al. Bayesian Physics Informed Neural Networks for real-world nonlinear dynamical systems. Comput. Methods Appl. Mech. Engrg. 402, 115346. issn: 0045-7825 (2022).

36. Saurabh, A., Fazel, M., Safar, M., Sgouralis, I. & Pressé, S. Single-photon smFRET. I: Theory and conceptual basis. en. Biophys. Rep. 3, 100089. issn: 2667-0747 (2023).

37. Saurabh, A., Safar, M., Fazel, M., Sgouralis, I. & Pressé, S. Single-photon smFRET: II. Application to continuous illumination. en. Biophys. Rep. 3, 100087. issn: 2667-0747 (2023).

38. Safar, M. et al. Single-photon smFRET. III. Application to pulsed illumination. en. Biophys. Rep. 2, 100088. issn: 2667-0747 (2022).

39. Gopich, I. V., Louis, J. M. & Chung, H. S. Maximum Likelihood Analysis of Diffusing Molecules with Conformational Dynamics in Single-Molecule FRET. J. Phys. Chem. B 129, 2187. issn: 1520-5207 (2025).

40. Jurich, C., Shao, Q., Ran, X. & Yang, Z. J. Physics-based modeling in the new era of enzyme engineering. Nat. Comput. Sci. 5, 279. issn: 2662-8457 (2025).

41. Fan, C. et al. A likelihood-based framework for demographic inference from genealogical trees. Nat. Genet. 57, 865. issn: 1546-1718 (2025).

42. Xu, L. W., Jazani, S., Kilic, Z. & Pressé, S. Single-molecule reaction-diffusion. Biophys. J. issn: 0006-3495 (2025).

43. Pressé, S. & Sgouralis, I. Data Modeling for the Sciences: Applications, Basics, Computations isbn: 9781009098502 (Cambridge University Press, 2023).

44. Sgouralis, I. et al. BNP-Track: a framework for superresolved tracking. Nat. Methods 21, 1716. issn: 1548-7105 (2024).

45. Swendsen, R. H. & Wang, J.-S. Replica Monte Carlo Simulation of Spin-Glasses. Phys. Rev. Lett. 57, 2607. issn: 0031-9007 (21 1986).

46. Earl, D. J. & Deem, M. W. Parallel tempering: Theory, applications, and new perspectives. Phys. Chem. Chem. Phys. 7, 3910. issn: 1463-9084 (23 2005).

47. Goodman, J. & Weare, J. Ensemble samplers with affine invariance. Commun. Appl. Math. Comput. Sci. 5, 65. issn: 1559-3940 (2010).

48. Wang, X. & Dunson, D. B. Parallelizing MCMC via Weierstrass Sampler 2014. 1312.4605 [stat.CO].

49. Wang, X., Guo, F., Heller, K. A. & Dunson, D. B. Parallelizing MCMC with Random Partition Trees in Advances in Neural Information Processing Systems (eds Cortes, C., Lawrence, N., Lee, D., Sugiyama, M. & Garnett, R.) 28 (Curran Associates, Inc., 2015). https://proceedings.neurips.cc/paper_files/paper/2015/file/40008b9a5380fcacce3976bf7c08af5b-Paper.pdf.

50. Scott, S. L. et al. Bayes and big data: the consensus Monte Carlo algorithm. Int. J. Manage. Sci. Eng. Manage. 11, 78. issn: 1750-9661 (2016).

51. Srivastava, S., Li, C. & Dunson, D. B. Scalable Bayes via Barycenter in Wasserstein Space. J. Mach. Learn. Res. 19, 1. http://jmlr.org/papers/v19/17-084.html (2018).

52. Nemeth, C. & Sherlock, C. Merging MCMC Subposteriors through Gaussian-Process Approximations. Bayesian Anal. 13. issn: 1936-0975 (2018).

53. Mesquita, D., Blomstedt, P. & Kaski, S. Embarrassingly Parallel MCMC using Deep Invertible Transformations in Proceedings of The 35th Uncertainty in Artificial Intelligence Conference (eds Adams, R.P. & Gogate, V.) 115 (PMLR, 2020), 1244. https://proceedings.mlr.press/v115/mesquita20a.html.

54. Jacob, P. E., O’Leary, J. & Atchadé, Y. F. Unbiased Markov Chain Monte Carlo Methods with Couplings. J. R. Stat. Soc. Ser. B Stat. Method. 82, 543. issn: 1467-9868. eprint: https://academic.oup.com/jrsssb/article-pdf/82/3/543/49324120/jrsssb_82_3_543.pdf (2020).

55. Glatt-Holtz, N. E., Holbrook, A. J., Krometis, J. A. & Mondaini, C. F. Parallel MCMC algorithms: theoretical foundations, algorithm design, case studies. Trans. Math. Appl. 8, tnae004. issn: 2398-4945. eprint: https://academic.oup.com/imatrm/article-pdf/8/2/tnae004/60767276/tnae004.pdf (2024).

56. Winter, S., Campbell, T., Lin, L., Srivastava, S. & Dunson, D. B. Emerging Directions in Bayesian Computation. Stat. Sci. 39, 62. issn: 0883-4237 (2024).

57. Sarmento, M. J. et al. The impact of the glycan headgroup on the nanoscopic segregation of gangliosides. Biophy. J. 120, 5530. issn: 0006-3495 (2021).

58. Davidović, D. et al. Which Moiety Drives Gangliosides to Form Nanodomains? J. Phys. Chem. Lett. 14, 5791. issn: 1948-7185 (2023).

59. Jonkman, J., Brown, C. M., Wright, G. D., Anderson, K. I. & North, A. J. Tutorial: guidance for quantitative confocal microscopy. Nat. Protoc. 15, 1585. issn: 1750-2799 (2020).

60. Galvanetto, N. et al. Extreme dynamics in a biomolecular condensate. Nature 619, 876. issn: 1476-4687 (2023).

61. Bi, G. et al. The ZAR1 resistosome is a calcium-permeable channel triggering plant immune signaling. Cell 184, 3528. issn: 0092-8674 (2021).

62. Ahn, J. H. et al. Phase separation drives aberrant chromatin looping and cancer development. Nature 595, 591. issn: 1476-4687 (2021).

63. Folkmann, A. W., Putnam, A., Lee, C. F. & Seydoux, G. Regulation of biomolecular condensates by interfacial protein clusters. Science 373, 1218. issn: 1095-9203 (2021).

64. Vaughan, J. C., Jia, S. & Zhuang, X. Ultrabright photoactivatable fluorophores created by reductive caging. Nat. Methods 9, 1181. issn: 1548-7105 (2012).

65. Victoria, A. H. & Maragatham, G. Automatic tuning of hyperparameters using Bayesian optimization. Evol. Syst. 12, 217. issn: 1868-6486 (2020).

66. Zhao, S., Li, K., Ahn, C. K., Huang, B. & Liu, F. Tuning-Free Bayesian Estimation Algorithms for Faulty Sensor Signals in State-Space. IEEE Trans. Ind. Electron. 70, 921. issn: 1557-9948 (2023).

67. Smith, C. S. et al. An automated Bayesian pipeline for rapid analysis of single-molecule binding data. Nat. Commun. 10. issn: 2041-1723 (2019).

68. Bryan IV, J. S., Sgouralis, I. & Pressé, S. Diffraction-limited molecular cluster quantification with Bayesian nonparametrics. Nat. Comput. Sci. 2, 102. issn: 2662-8457 (2022).

69. Kilic, Z., Schweiger, M., Moyer, C., Shepherd, D. & Pressé, S. Gene expression model inference from snapshot RNA data using Bayesian non-parametrics. en. Nat. Comput. Sci. 3, 174. issn: 2662-8457 (2023).

70. Chen, Z., Biggie, H., Ahmed, N., Julier, S. & Heckman, C. Kalman Filter Auto-Tuning With Consistent and Robust Bayesian Optimization. IEEE Trans. Aerosp. Electron. Syst. 60, 2236. issn: 2371-9877 (2024).

71. Saurabh, A. et al. Approaching maximum resolution in structured illumination microscopy via accurate noise modeling. npj Imaging 3. issn: 2948-197X (2025).

72. Gregg, C. & Hazelwood, K. Where is the data? Why you cannot debate CPU vs. GPU performance without the answer in (IEEE ISPASS) IEEE INTERNATIONAL SYMPOSIUM ON PERFORMANCE ANALYSIS OF SYSTEMS AND SOFTWARE (IEEE, 2011), 134.

73. Hennessy, J. L. & Patterson, D. A. Computer architecture. A quantitative approach Sixth edition (eds Patterson, D. A. & Asanović, K.) Literaturverzeichnis: Seite R-1-R-36, 936. isbn: 9780128119068 (Elsevier Science & Technology Books, Cambridge, Mass., 2017).

74. Beal, M., Ghahramani, Z. & Rasmussen, C. The Infinite Hidden Markov Model in Advances in Neural Information Processing Systems (eds Dietterich, T., Becker, S. & Ghahramani, Z.) 14 (MIT Press, 2001). https://proceedings.neurips.cc/paper_files/paper/2001/file/e3408432c1a48a52fb6c74d926b38886-Paper.pdf.

75. Teh, Y. W., Jordan, M. I., Beal, M. J. & Blei, D. M. Hierarchical Dirichlet Processes. J. Amer. Statistical Assoc. 101, 1566. issn: 1537-274X (2006).

76. Müller, P., Quintana, F. A., Jara, A. & Hanson, T. Bayesian Nonparametric Data Analysis isbn: 9783319368429 (Springer International Publishing AG, 2016).

77. Higham, N. J. & Mary, T. Mixed precision algorithms in numerical linear algebra. Acta Numer. 31, 347. issn: 1474-0508 (2022).

78. Micikevicius, P. et al. Mixed Precision Training 2018. 1710.03740 [cs.AI].

79. Wang, L. et al. Ladder: Enabling Efficient Low-Precision Deep Learning Computing through Hardware-aware Tensor Transformation in 18th USENIX Symposium on Operating Systems Design and Implementation (OSDI 24) (USENIX Association, Santa Clara, CA, 2024), 307. isbn: 978-1-939133-40-3. https://www.usenix.org/conference/osdi24/presentation/wang-lei.

80. Murray, R. et al. Randomized Numerical Linear Algebra: A Perspective on the Field With an Eye to Software 2023. 2302.11474 [math.NA].

81. Gabrié, M., Rotskoff, G. M. & Vanden-Eijnden, E. Adaptive Monte Carlo augmented with normalizing flows. Proc. Natl. Acad. Sci. U.S.A. 119, e2109420119. issn: 1091-6490 (2022).

82. Asghar, S., Pei, Q.-X., Volpe, G. & Ni, R. Efficient rare event sampling with unsupervised normalizing flows. Nat. Mach. Intell. 6, 1370. issn: 2522-5839 (2024).

83. Papamakarios, G., Nalisnick, E., Rezende, D. J., Mohamed, S. & Lakshminarayanan, B. Normalizing Flows for Probabilistic Modeling and Inference. Journal of Machine Learning Research 22, 1. http://jmlr.org/papers/v22/19-1028.html (2021).

84. Croitoru, F.-A., Hondru, V., Ionescu, R. T. & Shah, M. Diffusion Models in Vision: A Survey. IEEE Trans. Pattern Anal. Mach. Intell. 45, 10850. issn: 1939-3539 (2023).

85. Yang, L. et al. Diffusion Models: A Comprehensive Survey of Methods and Applications. ACM Comput. Surv. 56, 1. issn: 1557-7341 (2023).

86. Bezanson, J., Edelman, A., Karpinski, S. & Shah, V. B. Julia: A Fresh Approach to Numerical Computing. SIAM Rev. 59, 65. issn: 1095-7200 (2017).

87. Besard, T., Foket, C. & De Sutter, B. Effective Extensible Programming: Unleashing Julia on GPUs. IEEE Trans. Parallel Distrib. Syst. 30, 827. issn: 2161-9883 (2019).

88. Ansel, J. et al. PyTorch 2: Faster Machine Learning Through Dynamic Python Bytecode Transformation and Graph Compilation in Proceedings of the 29th ACM International Conference on Architectural Support for Programming Languages and Operating Systems, Volume 2 (ACM, 2024), 929.

89. Developers, T. TensorFlow 2025.

90. Innes, M. et al. Fashionable Modelling with Flux 2018.

91. Geman, S. & Geman, D. Stochastic Relaxation, Gibbs Distributions, and the Bayesian Restoration of Images. IEEE Trans. Pattern Anal. Mach. Intell. PAMI-6, 721. issn: 0162-8828 (1984).

92. Fluorescence Quantum Yields and Lifetimes for Alexa Fluor Dyes Table https://www.thermofisher.com/us/en/home/references/molecular-probes-the-handbook/tables/fluorescence-quantum-yields-and-lifetimes-for-alexa-fluor-dyes.html.

93. Innes, M. Flux: Elegant machine learning with Julia. J. Open Source Softw. 3, 602. issn: 2475-9066 (2018).

94. Dongarra, J., Hammarling, S., Higham, N. J., Relton, S. D. & Zounon, M. in EuroPar 2017: Parallel Processing 511 (Springer International Publishing, 2017). isbn: 9783319642031.

95. Dongarra, J. et al. The Design and Performance of Batched BLAS on Modern High-Performance Computing Systems. Procedia Comput. Sci. 108, 495. issn: 1877-0509 (2017).

96. Abdelfattah, A. et al. A Set of Batched Basic Linear Algebra Subprograms and LAPACK Routines. ACM Trans. Math. Softw. 47, 1. issn: 1557-7295 (2021).

97. Quinn, M. J. Parallel computing. Theory and practice 2., [rev.] ed., internat. ed., [Nachdr.] Literaturverz. S. 391–433. 446 pp. isbn: 0070512949 (McGraw-Hill, New York [u.a.], 1995).

98. Valiant, L. G. A bridging model for parallel computation. Commun. ACM 33, 103. issn: 1557-7317 (1990).

99. Introduction to parallel computing 2. ed., [reprint.] (ed Grama, A.) Includes bibliographical references and index. 636 pp. isbn: 9780201648652 (Pearson, Harlow [u.a.], 2011).

100. Pearl, J. Probabilistic Reasoning in Intelligent Systems. Networks of Plausible Inference Rev. 2. print. 584 pp. isbn: 1558604790 (Elsevier Science, 2014).

101. Drepper, U. What every programmer should know about memory https://people.freebsd.org/~lstewart/articles/cpumemory.pdf (2025).

102. Goodrich, M. T., Tamassia, R. & Mount, D. M. Data structures and algorithms in C++ 2. ed. Literaturverz. S. [697] - 701. 714 pp. isbn: 9780470383278 (Wiley, Hoboken, NJ, 2011).

103. Jones, R., Hosking, A. & Moss, E. The Garbage Collection Handbook: The Art of Automatic Memory Management isbn: 9781003276142 (Chapman and Hall/CRC, 2023).

104. Danisch, S. & Krumbiegel, J. Makie.jl: Flexible high-performance data visualization for Julia. J. Open Source Softw. 6, 3349. issn: 2475-9066 (2021).

105. Besançon, M. et al. Distributions.jl: Definition and Modeling of Probability Distributions in the JuliaStats Ecosystem. J. Stat. Softw. 98. issn: 1548-7660 (2021).

